# An open resource for nonhuman primate imaging

**DOI:** 10.1101/227462

**Authors:** Michael P. Milham, Lei Ai, Bonhwang Koo, Ting Xu, Fabien Balezeau, Mark G. Baxter, Paula L. Croxson, Christienne G. Damatac, Noam Harel, Winrich Freiwald, Timothy D. Griffiths, Stefan Everling, Benjamin Jung, Sabine Kastner, David A. Leopold, Rogier B. Mars, Ravi S. Menon, Adam Messinger, John H. Morrison, Jennifer Nacef, Jamie Nagy, Michael Ortiz Rios, Christopher I. Petkov, Mark Pinsk, Colline Poirier, Reza Rajimehr, Matthew F.S Rushworth, Brian E. Russ, Michael Schmid, Caspar M. Schwiedrzik, Jerome Sallet, Jakob Seidlitz, Leslie Ungerleider, Alexander Thiele, Doris Tsao, Essa Yacoub, Frank Ye, Wilbert Zarco, Daniel S. Margulies, Charles Schroeder

## Abstract

Non-human primate neuroimaging is a rapidly growing area of research that promises to transform and scale translational and cross-species comparative neuroscience.

Unfortunately, the technological and methodological advances of the past two decades have outpaced the accrual of data, which is particularly challenging given the relatively few centers that have the necessary facilities and capabilities. The PRIMate Data Exchange (PRIME-DE) addresses this challenge by aggregating independently acquired non-human primate magnetic resonance imaging (MRI) datasets and openly sharing them via the International Neuroimaging Data-sharing Initiative (INDI). Here, we present the rationale, design and procedures for the PRIME-DE consortium, as well as the initial release, consisting of 13 independent data collections aggregated across 11 sites (total = 98 macaque monkeys). We also outline the unique pitfalls and challenges that should be considered in the analysis of the non-human primate MRI datasets, including providing automated quality assessment of the contributed datasets.

## BACKGROUND AND SUMMARY

Translational and cross-species comparative neuroscience research enables a bridging of knowledge across both invasive and noninvasive approaches. A growing body of research has documented the utility of magnetic resonance imaging (MRI) technologies to support in vivo examination of brain organization and function in non-human primates (Vanduffel W n.d.; Rilling 2014; D. C. Van Essen and Glasser n.d.; Zhang D n.d.; Shmuel and Leopold n.d.). Recent work has demonstrated the ability to recapitulate findings from gold-standard invasive methodologies (Ghahremani et al. 2017); (Donahue et al. 2016), as well as provide novel insights into the organizational principles of the non-human primate connectome (Goulas et al. 2017; Hutchison and Everling 2014; Hutchison et al. 2011; Vincent et al. 2007) and cross-species comparative connectomics (Hutchison et al. 2015; Miranda-Dominguez et al. 2014; Hutchison et al. 2012; Mars et al. 2011)(Seidlitz, Váša, et al. 2017), which could only be afforded through in vivo studies. These advances are timely given the growing prominence of large-scale national and international initiatives focused on advancing our understanding of human brain organization and the ability to generate novel therapeutics for neurology and psychiatry (Bargmann and Newsome 2014).

Despite the various demonstrations of feasibility and utility, the field of non-human primate neuroimaging is still in its early stages. Numerous unique challenges related to the acquisition and processing of non-human primate data are still being addressed (e.g., (Seidlitz, Sponheim, et al. 2017; R. Matthew Hutchison 2012)), and the potential for broad reaching cross-species studies remains to be explored. We introduce the Primate Data Exchange (PRIME-DE) to create an open science resource for the neuroimaging community that will facilitate the mapping of the non-human primate connectome. To accomplish this, we aggregate a combination of anatomical, functional, and diffusion MRI datasets from laboratories throughout the world, and make these data available to the scientific community.

## METHODS

### Criteria for data contributions

PRIME-DE welcomes contributions from any laboratory willing to openly share multimodal MRI datasets obtained from non-human primates, including but not limited to functional MRI, diffusion MRI and structural MRI. Contributors are responsible for ensuring that any data collected and shared were obtained in accordance with local ethical and regulatory requirements.

There are no set exclusion criteria. We encourage the sharing of all data, independent of quality. This decision is based on the realizations that: 1) there is no consensus on acceptable criteria for movement in functional MRI or diffusion MRI data, 2) high motion datasets are essential to the determination of the impact of motion on reliability, and 3) new approaches continue to be developed to account for movement artifacts. We also encourage submission of data from other modalities (e.g., ASL) or experimental paradigms (e.g. longitudinal data, pharmacologic manipulations) when available.

### Data preparation and aggregation

PRIME-DE data aggregation is carried out through the International Neuroimaging Data-sharing Initiative (INDI)(Mennes et al. 2013) portal located at the Neuroimaging Informatics Tools and Resources Clearinghouse (NITRC) (http://fcon_1000.projects.nitrc.org/indi/indiPRIME.html). Following the model of prior efforts, all contributions are reviewed by the INDI team following upload and corrected as needed to ensure consistent data organization within and across sites. Before open release, each contributing site reviews their reorganized phenotypic records, five random images per imaging modality and their collection-specific narrative for final approval.

## DATA RECORDS

### Overview

At present, PRIME-DE contains 13 collections aggregated across 11 sites; data from a total of 98 monkeys is included to date (See Table 1 for information on each institution). To promote usage of a standardized data format, all data are organized using the Brain Imaging Data Structure (BIDS) format. All PRIME-DE datasets can be accessed through the PRIME-DE site (http://fcon_1000.projects.nitrc.org/indi/indiPRIME.html). Prior to downloading the data, users are required to establish a user account on NITRC and register with INDI (anticipated time: < 1 minute).

**Table 1.**
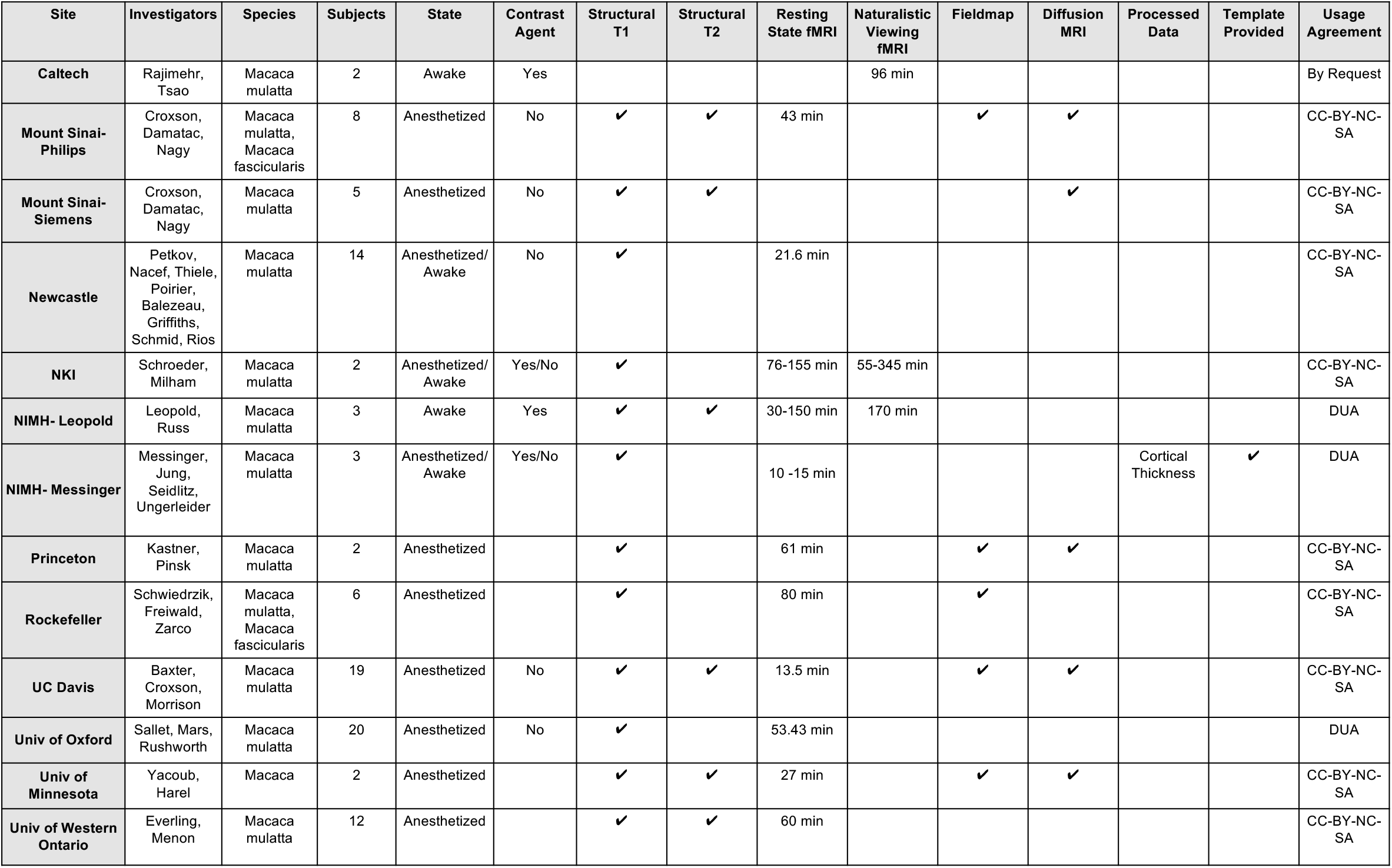
Experimental Design

### Phenotypic information

Given that this is a retrospective data collection, we focus on basic phenotypic measures that are relatively standard in the neuroimaging field, as well as those fundamental for analyses and sample characterization. Minimal phenotypic information includes: age, sex, species. The contribution of additional variables that can enhance data usage is encouraged, though not required.

### MRI data

For each of the 11 PRIME-DE collections, for each unique ID #, at least one structural MRI (sMRI), and one corresponding R-fMRI dataset are available. In addition, one collection (NIMH) also provided cortical thickness data and R-fMRI data aligned to an anatomical template. Corresponding diffusion MRI (dMRI) datasets are available for five collections. Fieldmap images for fMRI correction are available for five collections. Consistent with its popularity in the imaging community and prior usage in INDI efforts, the NIFTI file format was selected for storage of the PRIME-DE MRI datasets. Table 2 lists the specific MRI scanners and head coils utilized for each collection. Specific MRI sequence parameters for the various data collections are summarized in Tables 2, 3, 4 and detailed on the PRIME website. Across collections, R-fMRI acquisition durations varied from 8 to 345 minutes per subject; in two collections, subjects were in an awake state; in three collections, subjects were scanned both awake and under anesthesia; in the remaining eight collections, subjects were scanned under anesthesia. Along with R-fMRI, two collections provided Naturalistic Viewing fMRI. **3.3.** Data Licensing Contributors to PRIME will be able to set the sharing policy for their data in accord with their preferences and institutional requirements. For each sample, the contributor will set the sharing permissions for their data using one or more the following four policies:

**Table 2.** Scanner Information

| Site | Manufacturer | Model | Field Strength (T) | Head coil # channels |
| --- | --- | --- | --- | --- |
| Caltech | Siemens | Tim Trio | 3 | 8 |
| Mount Sinai- Philips | Philips | Achieva | 3 | 4 |
| Mount Sinai- Siemens | Siemens | Skyra | 3 | 8 |
| Newcastle | Bruker | Vertical Bruker | 4.7 | 4-8 |
| NKI | Siemens | Tim Trio | 3 | 8 |
| NIMH- Leopold | Bruker | Biospec Vertical | 4.7 | 8 |
| NIMH- Messinger | Bruker | Biospec Vertical | 4.7 | 1-4 |
| Princeton | Siemens | Prisma VD11C | 3 | Siemens Loop Coil, Large (11cm) |
| Rockefeller | Siemens | TIM Trio + AC88 gradient | 3 | 8-channel phased-array receive with a single loop transmit |
| UC Davis | Siemens | Skyra | 3 | 4 |
| Univ of Oxford |  |  | 3 | 4 |
| Univ of Minnesota | Siemens | Syngo B17 | 7 | 16 channel transmit/receive + 6 receive only |
| Univ of Western Ontario | Siemens 7T | Magnetom | 7 |  |

**Table 3.**
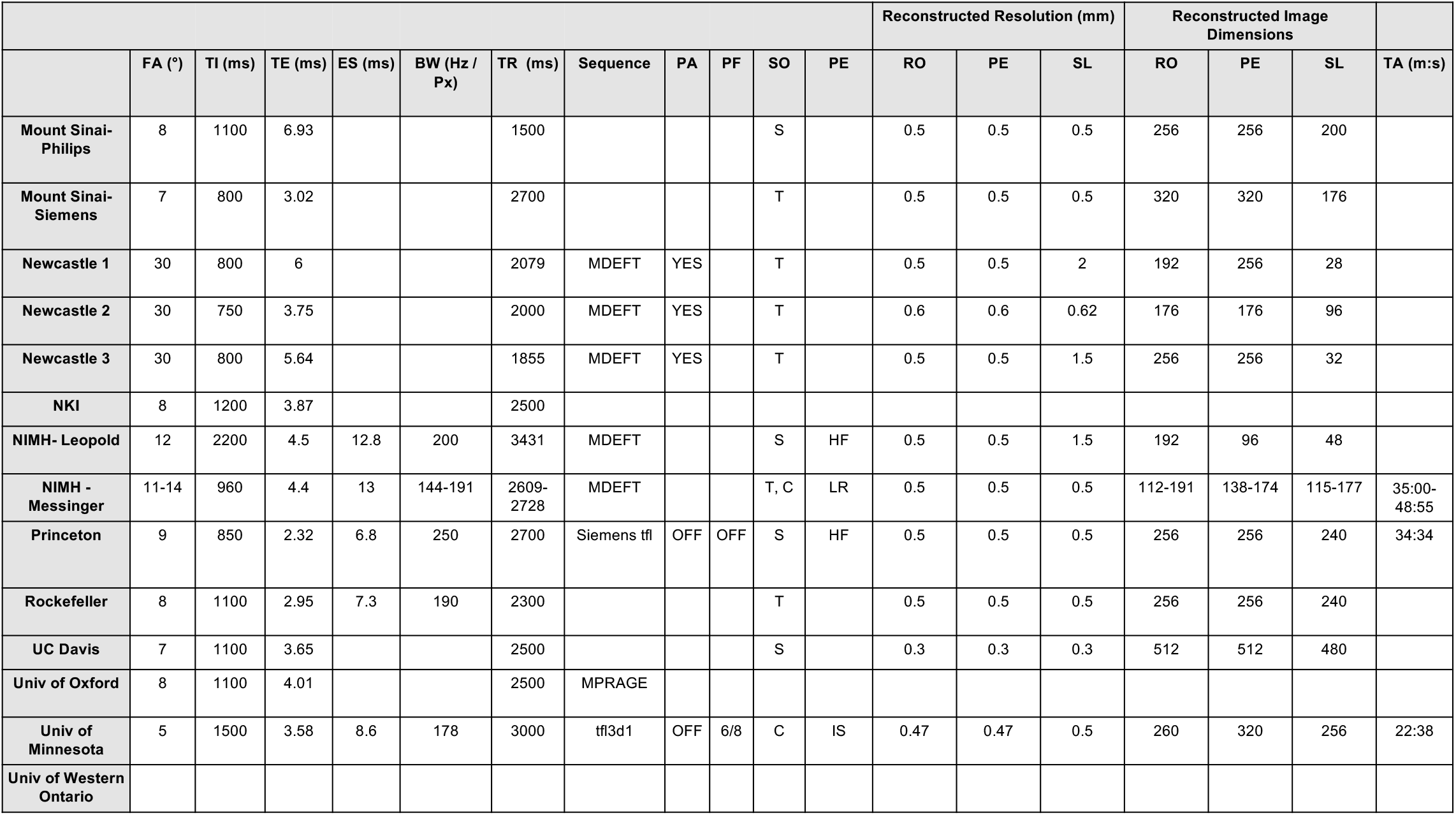
Structural MRI Sequence Information

**Table 4.**
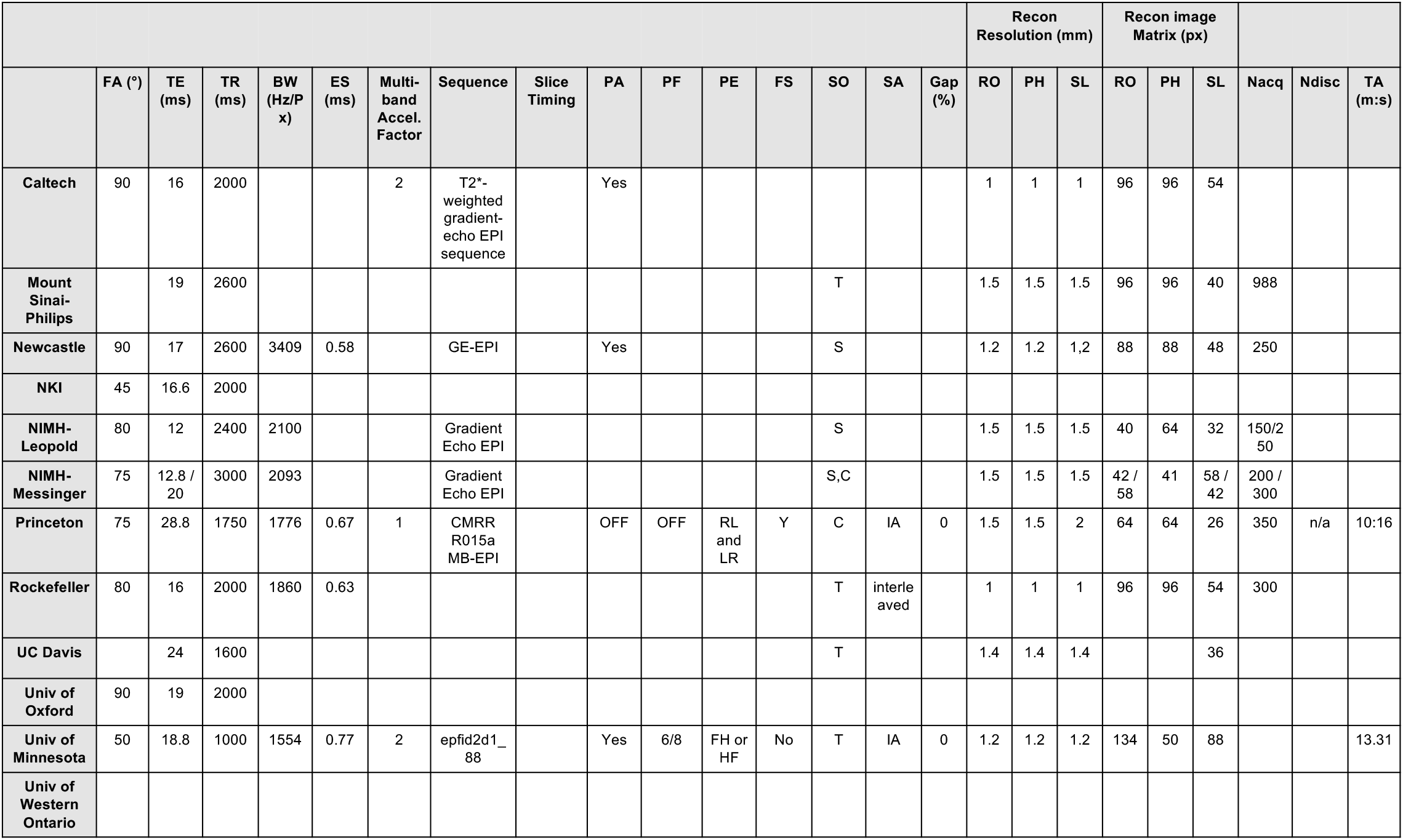
Functional MRI Sequence Information

1. *Creative Commons – Attribution-NonCommercial Share Alike (CC-BY-NC-SA)*. https://creativecommons.org/licenses/by-nc-sa/4.0/ Standard INDI data sharing policy. Prohibits use of the data for commercial purposes.
2. *Creative Commons – Attribution (CC-BY)*. https://creativecommons.org/licenses/by/4.0/ Least restrictive data sharing policy.
3. *Custom Data Usage Agreement*. Users must complete a data usage agreement (DUA) prior to gaining access to the data. Contributors can customize the agreement as they see fit, including determining whether or not signatures from authorized institutional official are required prior to executing the DUA.
4. *By Request* Most Restrictive data sharing policy; requires contact of the investigator and likely formal collaboration to obtain data

## TECHNICAL VALIDATION

### 4.1 Automated Quality Assessment

Consistent with the established policy of INDI, all data contributed to PRIME was made available to users regardless of data quality. The rationale of this decision has been the lack of consensus on optimal quality criteria in regards to specific measures or their combinations and cutoffs - a reality that is even more pronounced in nonhuman primate imaging given the variation in data quality and characteristics across scan protocols given the lack of harmonization.

Following the tradition of recent INDI data-sharing consortia, a collection of automated, reference-free quality assurance measures, known as the Preprocessed Connectome Project Quality Assurance Protocol (PCP-QAP), is being made available with the PRIME datasets. These measures focus on structural and temporal (when appropriate) aspects of the datasets. Table 5 provides a brief description of the measures included, and Figures 1 and 2 depict a subset of QAP results(Magnotta, Friedman, and FIRST BIRN 2006; Mortamet et al. 2009; Giannelli et al. 2010; Jenkinson et al. 2002). As would be expected, measures of head motion are notably smaller for sites using anesthetized scan sessions than awake (NIMH-Russ/Leopold, NIMH-Messinger, NKI, Newcastle).

**Table 5.** Description of PCP QAP Measures

| Spatial Metrics | Description |
| --- | --- |
| Contrast-to-noise ratio (CNR) (Magnotta et al. 2006) (sMRI only) | $M^{GM} \text{ intensity} - M^{WM} \text{ intensity} / SD^{air} \text{ intensity}$. Larger values reflect a better distinction between WM and GM. |
| Artifactual voxel detection (Qi1) (Mortamet et al. 2009) (sMRI only) | voxels with intensity corrupted by artifacts/voxels in the background. Larger values reflect more artifacts which likley due to motion or image instability. |
| Smoothness of Voxels (FWHM) (Friedman et al. 2006) † | Full-width half maximum of the spatial distribution of the image intensity values. Larger values reflect more spatial smoothing perhaps due to motion or technical differences. |
| Signal-to-noise ratio (SNR) (Magnotta et al. 2006) | $M^{GM} \text{ intensity} / SD^{air} \text{ intensity}$. Larger values reflect less noise. |
| Temporal Metrics (fMRI and DTI only) | Description |
| Ghost to Signal Ratio (GSR) (Gianelli et al. 2010) † | M signal in the 'ghost' image divided by the M signal within the brain. Larger values reflect more ghosting likely due to physiological noise, motion, or technical issues. |
| Mean framewise displacement- Jenkinson (meanFD) (Jenkinson et al. 2002) ‡ | Sum absolute displacement changes in the x, y and z directions and rotational changes around them. Rotational changes are given distance values based on changes across the surface of a 50 mm radius sphere. Larger values reflect more movement. |
| Standardized DVARS (Nichols 2012) ‡ | Spatial SD of the data temporal derivative normalized by the temporal SD and autocorrelation. Larger values reflect larger frame-to-frame differences in signal intensity due to head motion or scanner instability. |
| Global Correlation (GCORR) ‡ | M correlation of all combinations of voxels in a time series. Illustrates differences between data due to motion/physiological noise. Larger values reflect a greater degree of spatial correlation between slices, which may be due to head motion or 'signal leakage' in simultaneous multi-slice acquisitions. |

† For R-fMRI data these metrics are computed on mean functional data.
‡ For R-fMRI these metrics are computed on time series data. M, Mean; GM, Gray Matter; WM, White Matter; s.d., Standard Deviation.
**Adopted from:** Di Martino A, et al. 2017. Enhancing studies of the connectome in autism using the autism brain imaging data exchange II. Sci Data. 4:170010.

**Figure 1.**
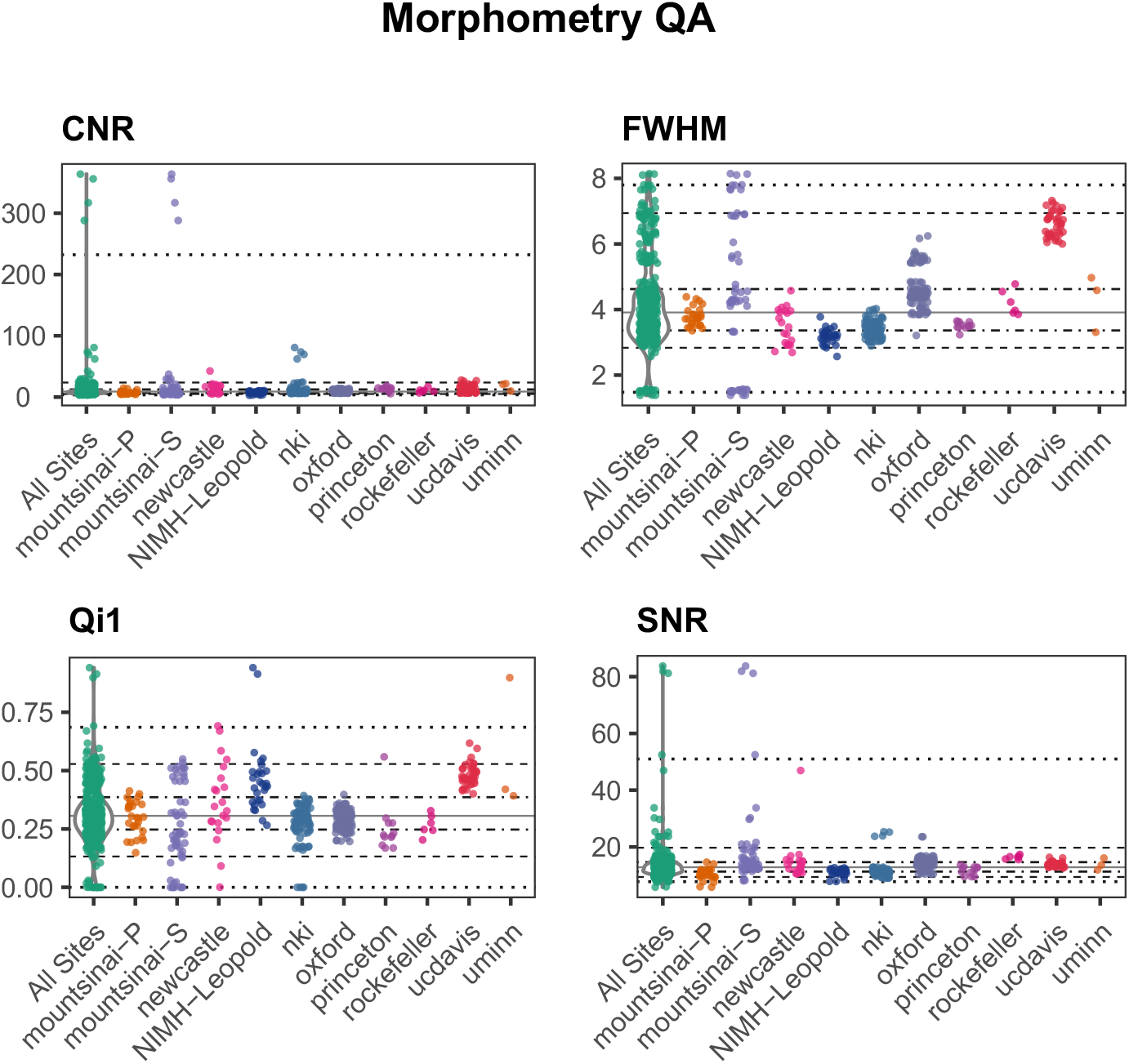
Spatial quality metrics for morphometry MRI datasets

**Figure 2.**
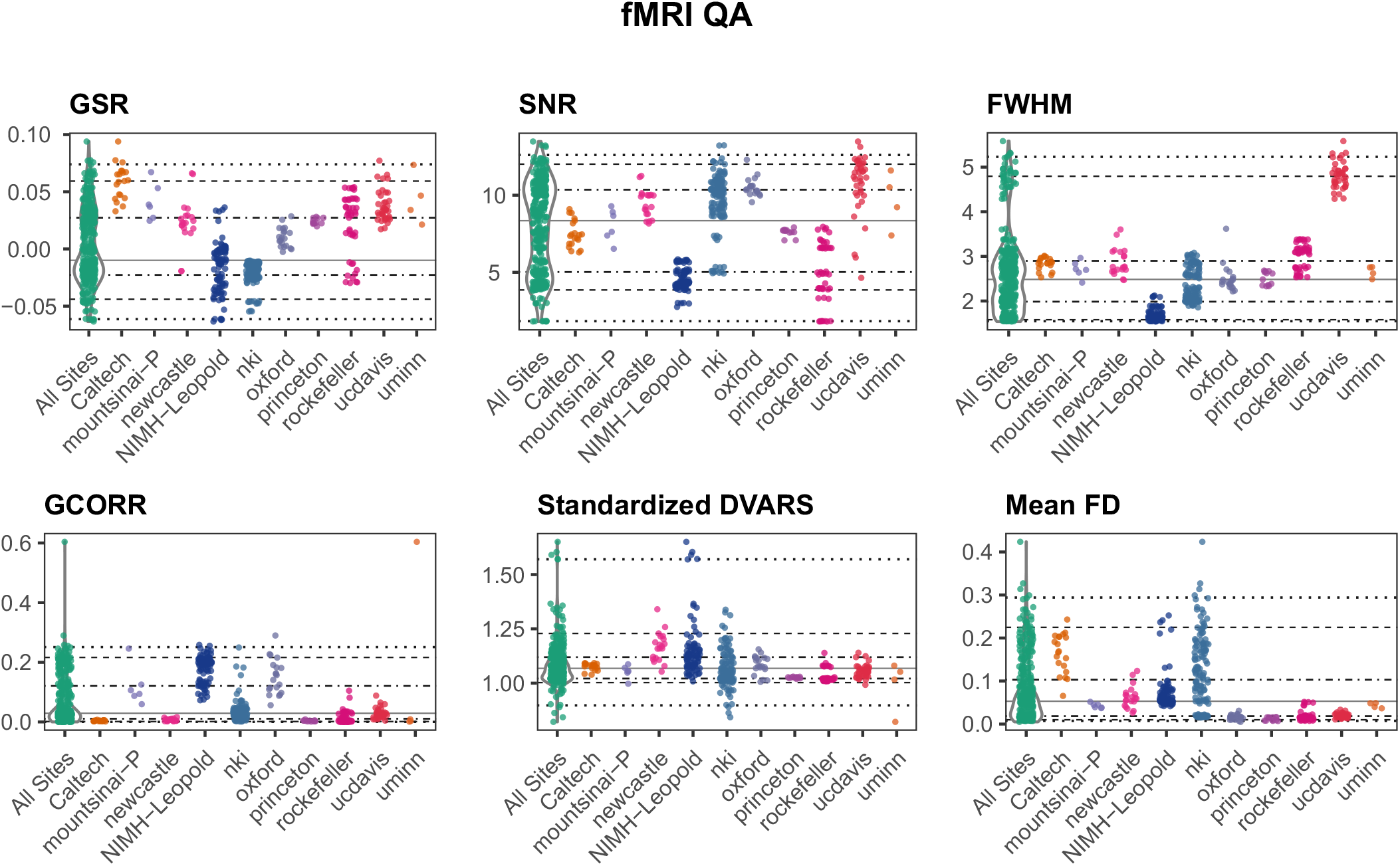
Spatial and temporal quality metrics for functional MRI (fMRI)

## USAGE NOTES

### 5.1 Challenges in the Processing of Nonhuman Primate Imaging data

There are a variety of challenges faced when trying to adapt well-established methods for human neuroimaging processing to monkey and rodent data. Beyond the differences between species in tissue contrast, brain shape and size, and type and amount of tissue surrounding the brain, there are significant differences in data collection equipment and acquisition protocols. Non-human primate data are often acquired at very high fields (4.7T, 7T, 9.4T, 11.7T), using some non-standardized arrangement of surface coils. These result in increased variations in image intensity due to B1 inhomogeneity and non-uniform coil coverage, and greater distortion and dephasing due to susceptibility. Another issue is that the equipment and acquisition protocols used are typically customized, resulting in substantial variation in the quality and characteristics of data collected at different sites. Consequently, there is no one-size fits all strategy for processing animal data and researchers need a good deal of flexibility to optimize their pipelines for the data at hand.

Brain extraction and tissue segmentation are more challenging in non-human imaging data due to differences in tissue contrast and the nature of structures immediately surrounding the brain. If compromised, these steps in turn can dramatically compromise image registration and normalization procedures, as well as temporal denoising approaches. As of yet, there is no consensus optimal solution for each of these processing steps, in part due to the many sources of variation across studies that can differentially impact data characteristics and quality (e.g., anesthesia protocols, coil type, use of contrast agents, magnet strength, animal/rodent type). Additionally, commonly used pre-processing pipelines, used extensively with human neuroimaging datasets, often fail to work properly on non-human primate datasets. As a result, researchers commonly work to optimize individual steps for their datasets outside of traditional workflows, resulting in different pipelines and processing steps across groups. There are efforts underway to form best practices to guide this process and help researchers avoid the need to redefine pipelines themselves (e.g., (Seidlitz, Sponheim, et al. 2017; “The Average Baboon Brain: MRI Templates and Tissue Probability Maps from 89 Individuals” 2016)), however currently it is still necessary for researchers to do so.

### 5.2 Resources and Solutions

#### 5.2.1. Templates and Atlases

A number of macaque templates were created in the last decade, including single animal templates e.g. the NeuroMap macaque atlas (Dubach and Bowden 2009) and the 3D Digital D99 Template (Reveley et al. 2017), and population-averaged templates based on multiple animals e.g. 112RM-SL (McLaren et al. 2009), INIA19 (Integrative Neuroscience Initiative on Alcoholism, (Rohlfing et al. 2012)), MNI (Montreal Neurological Institute, (Frey et al. 2011)), and the most recent NMT (National Institute of Mental Health Macaque Template, (Seidlitz, Sponheim, et al. 2017)). In addition, there are surface-based atlases, including the macaque single-subject F99 atlas (David C. Van Essen 2012, 2002) and the group-average Yerkes19 macaque atlas (Donahue et al. 2016). Data collected in individual macaques can be aligned to these templates using affine and non-linear registration. These templates provide a common anatomical space and coordinate system for specifying specific brain locations and visualizing data collected across days, animals, and laboratories.

Some of these templates link to volumetric digital brain atlases ((Frey et al. 2011); (Reveley et al. 2017)(Seidlitz, Sponheim, et al. 2017)(Reveley et al. 2017; Saleem and Logothetis 2012; Seidlitz, Sponheim, et al. 2017) derived from analysis of histological tissue (Saleem and Logothetis 2012; Paxinos, Huang, and Toga 1999; Paxinos 2009). These anatomical parcellations can be warped to individual subjects using standard linear and non-linear registration algorithms (e.g., AFNI’s 3dAllineate and 3dQwarp). Scripts to automate this alignment are available for the single-subject D99 template (http://afni.nimh.nih.gov/pub/dist/atlases/macaque), and the recently published National Institute of Mental Health Macaque Template (NMT;(Seidlitz, Sponheim, et al. 2017)) (https://afni.nimh.nih.gov/NMT). The NMT is a high resolution (0.25 mm isotropic) T1 template built from *in vivo* scans of 31 young adult macaques. This volume (and accompanying surfaces) is representative of the adult population and provides anatomical detail akin to that of *ex vivo* templates, which require days of scanning to acquire. The NMT is available via the PRIME-DE website as well as on GitHub (https://github.com/ims290/NMT). The database also includes resting state data from 3 subjects that have been aligned to the NMT (see NIMH-Messinger in Table 1). A similar multi-subject template also exists for pre-pubertal rhesus monkeys (Fox et. al, 2015).

Other anatomical parcellations have been defined on the surface using the single-subject F99 template (available in Caret; (David C. Van Essen et al. 2012)), which can be used for analysis on the cortical sheet. For example, the cortical parcellation from (Markov et al. 2014) includes quantitative tract-tracing connectivity estimates for a subset of these regions.

#### 5.2.2. Improving skull extraction, segmentation and registration

A high quality T1 image with isotropic voxels is important for skull extraction. There are a number of brain extraction algorithms and available tools, e.g. the Brain Extraction Tool (BET in FSL), 3dSkullStip in AFNI, the Hybrid Watershed Algorithm (HWA in FreeSurfer), BSE in BrainSuite, Robust Brain Extraction (BOBEX), and ANTs. Most of these tools do a good job for human data, however, the performance is suboptimal and variable in NHP due to the differences in brain structure (e.g. size, adipose tissue, olfactory bulb) and the quality of the T1 image (SNR, inhomogeneous intensity). Accordingly, the parameters and/or related atlas library need to be customized to optimize the brain extraction in NHP. For example, in AFNI the program “3dSkullStrip” with alternative options “–monkey” “-marmoset” and “–surface_coil” are available for brain extraction in NHP. Population brain templates, such as the NIMH Macaque Template (NMT), can further improve and automate the registration and brain extraction process (Seidlitz, Sponheim, et al. 2017).

Standard segmentation algorithms can separate gray versus white matter but if the signal is not homogenous, which is typically the case at higher magnetic fields, segmentation in some parts of the brain will be better than others (especially subcortically). Registration of T2 datasets to T1 structural scans also remains a challenge. Affine or non-linear registration algorithms can work well provided that intermediate scans are available. For instance, a full brain T1 structural scan from the same monkey obtained along with T2 images (also with as much coverage of the brain as possible) could be crucial for registering T2 datasets to any of the freely available monkey template brains that are registered to macaque atlases.

One way to reduce or eliminate the manual intervention during brain extraction and tissue segmentation - using only the typically-acquired T1 scan - is to rely on priors defined on a high-resolution and high-contrast template. The multi-subject NMT includes manually refined masks of the brain, cortical gray matter, and various tissue types (including blood vasculature) (Seidlitz, Sponheim, et al. 2017). Applying the inverse anatomical alignment transformations to the NMT brain mask produces an approximate single subject mask for brain extraction. A more precise individual brain mask and tissue segmentation can be obtained using the NMT’s representative brain and tissue segmentation masks as priors.

The NMT distribution includes scripts that use AFNI and ANTs to perform these mask refinements (as well as morphological analysis). These improvements could be critical for later processing steps for functional MRI data. Furthermore, the NMT includes surfaces for visualization of individual subject or group results in a standard coordinate space. Future work could add to these advances, such as tailoring existing surface-based processing pipelines (e.g., CIVET or FreeSurfer) to be specifically used with non-human primate MRI data.

#### 5.2.3. Head Motion

Head motion in NHP imaging is an important concern, just as it is in human neuroimaging studies. For the most part, one can apply human imaging motion correction techniques to NHP data directly. However there are a few concerns with NHP neuroimaging that will be addressed below.

Anesthesia is commonly used in NHP functional neuroimaging, in part due to the lower behavioral and technical demands than required to achieve awake imaging. As reflected by the QAP results, another benefit is that anesthesia dramatically reduces motion artifacts during NHP scanning. However, the use of anesthesia comes with its own set of tradeoffs dealing with how the drugs used interact with neural activity. There are changes in FC patterns based on the particular set and doses of agents used, and in comparison to awake imaging(Xu et al. 2017). For this reason, researchers should always assess how anesthesia may, or may not, influence the results of their study before using it.

For awake NHP imaging, the animals are far more likely to create motion artifacts that need to be addressed during preprocessing and subsequent analyses. Proper training and acclimation to the chair and scanner setup are of great importance in reducing the amount of head motion. As with human neuroimaging best practices, keeping individual scan periods to the shortest necessary for your task will help to reduce motion artifacts. Recent human studies also suggested that movies (naturalistic viewing) paradigm may help to reduce head motion relative to resting conditions (e.g., (Vanderwal et al. 2015)(Alexander et al. 2017). This is also true in awake NHP imaging; for example in PRIME-NKI site, the mean FD for rest sessions was 0.21 (SD=0.03) but 0.14 (SD=0.07) during movie sessions (t=2.82, p=0.006).

Regarding motion correction algorithms,those designed for human neuroimaging data similarly for NHP data. As such, most groups use SPM, AFNI, ANTs, or FSL software to estimate the motion parameters and remove motion artifacts. The estimates of the movement values can be used as regressors of no interest during the analysis of functional data, if desired. The grayplot proposed by (Power 2017) can be used to illustrate the motion and the denoising effects. However, as with all neuroimaging data, image distortions or signal drop-out with caused movement correction to be suboptimal.

## AVAILABILITY OF SUPPORTING DATA

### LIST OF ABBREVIATIONS

dMRI: : diffusion magnetic resonance imaging
DUA: : data usage agreement
fMRI: : functional magnetic resonance imaging
INDI: : International Neuroimaging Data-sharing Initiative
MRI: : magnetic resonance imaging
NIMH: : National Institute of Mental Health
NIH: : National Institutes of Health
NMT: : National Institute of Mental Health Macaque Template
QA: : quality assurance
QAP: : quality assurance protocol
sMRI: : structural magnetic resonance imaging
FA: : flip angle
TE: : echo time
TR: : repetition time
BW: : bandwidth per pixel
ES: : echo spacing
PA: : parallel acquisition
PF: : Partial Fourier (half scan)
PE: : Phase encoding
FS: : fat suppression
SO: : slice orientation
SA: : slice acquisition order
Gap: : gap between slices
SO: : Slice orientation
PE: : phase encoding
RO: : read out direction
Nacq: : number of volumes collected
Ndisc: : number of initial volumes discarded by the scanner
TA: : acquisition time

## ETHICS APPROVAL AND CONSENT TO PARTICIPATE

All experimental procedures were approved by local ethics boards prior to any data collection. UK macaque datasets were obtained with Home Office approval and abide with the European Directive on the protection of animals used in research (2010/63/EU).

## CONSENT FOR PUBLICATION

Not appropriate

## AUTHOR CONTRIBUTIONS

### Conception and Experimental Design

DSM, MPM, CS

### Implementation and Logistics

LA, DSM, MPM, CS, BK, TX

### Data Collection

MPM, FB, MGB, PLC, CGD, NH, WF, TDG, BJ, SK, DAL, RBM, AM, JHM, JNac, JNag, MOR,CIP, MP, CP, RR, MSFR, BER, MS, CMS, JSa, JSe, LU, AT, DT, EY, FY, WZ, DSM, CES

### Data Informatics

BK, LA

### Data Analysis

LA, MPM

### Drafting of the Manuscript

MPM

### Critical Review and Editing of the Manuscript

All authors contributed to the critical review and editing of the manuscript.

## ACKNOWLEDGEMENTS

We thank Cameron Craddock for his creation and open sharing of the Preprocessed Connectome Project (PCP) Quality Assessment Project (QAP), which was used here. (http://preprocessed-connectomes-project.org/quality-assessment-protocol/index.html).

## REFERENCES

Alexander, Lindsay M., Jasmine Escalera, Lei Ai, Charissa Andreotti, Karina Febre, Alex Mangone, Natan Vega Potler, et al. 2017. “An Open Resource for Transdiagnostic Research in Pediatric Mental Health and Learning Disorders.” bioRxiv. https://doi.org/10.1101/149369.

Bargmann, Cornelia I., and William T. Newsome. 2014. “The Brain Research Through Advancing Innovative Neurotechnologies (BRAIN) Initiative and Neurology.” JAMA Neurology 71 (6). American Medical Association:675–76.

Donahue, Chad J., Stamatios N. Sotiropoulos, Saad Jbabdi, Moises Hernandez-Fernandez, Timothy E. Behrens, Tim B. Dyrby, Timothy Coalson, et al. 2016. “Using Diffusion Tractography to Predict Cortical Connection Strength and Distance: A Quantitative Comparison with Tracers in the Monkey.” The Journal of Neuroscience: The Official Journal of the Society for Neuroscience 36 (25):6758–70.

Dubach, Mark F., and Douglas M. Bowden. 2009. “BrainInfo Online 3D Macaque Brain Atlas: A Database in the Shape of a Brain.” presented at the Society of Neuroscience Annual Meeting, Chicago, IL.

Frey, Stephen, Deepak N. Pandya, M. Mallar Chakravarty, Lara Bailey, Michael Petrides, and D. Louis Collins. 2011. “An MRI Based Average Macaque Monkey Stereotaxic Atlas and Space (MNI Monkey Space).” NeuroImage 55 (4):1435–42.

Ghahremani, Maryam, R. Matthew Hutchison, Ravi S. Menon, and Stefan Everling. 2017. “Frontoparietal Functional Connectivity in the Common Marmoset.” Cerebral Cortex 27 (8):3890–3905.

Giannelli, Marco, Stefano Diciotti, Carlo Tessa, and Mario Mascalchi. 2010. “Characterization of Nyquist Ghost in EPI-fMRI Acquisition Sequences Implemented on Two Clinical 1.5 T MR Scanner Systems: Effect of Readout Bandwidth and Echo Spacing.” Journal of Applied Clinical Medical Physics / American College of Medical Physics 11 (4):3237.

Goulas, Alexandros, Peter Stiers, R. Matthew Hutchison, Stefan Everling, Michael Petrides, and Daniel S. Margulies. 2017. “Intrinsic Functional Architecture of the Macaque Dorsal and Ventral Lateral Frontal Cortex.” Journal of Neurophysiology 117 (3):1084–99.

Hutchison, R. Matthew, Jody C. Culham, J. Randall Flanagan, Stefan Everling, and Jason P. Gallivan. 2015. “Functional Subdivisions of Medial Parieto-Occipital Cortex in Humans and Nonhuman Primates Using Resting-State fMRI.” NeuroImage 116 (August):10–29.

Hutchison, R. Matthew, and Stefan Everling. 2014. “Broad Intrinsic Functional Connectivity Boundaries of the Macaque Prefrontal Cortex.” NeuroImage 88 (March):202–11.

Hutchison, R. Matthew, Jason P. Gallivan, Jody C. Culham, Joseph S. Gati, Ravi S. Menon, and Stefan Everling. 2012. “Functional Connectivity of the Frontal Eye Fields in Humans and Macaque Monkeys Investigated with Resting-State fMRI.” Journal of Neurophysiology 107 (9):2463–74.

Hutchison, R. Matthew, L. Stan Leung, Seyed M. Mirsattari, Joseph S. Gati, Ravi S. Menon, and Stefan Everling. 2011. “Resting-State Networks in the Macaque at 7 T.” NeuroImage 56 (3):1546–55.

Jenkinson, Mark, Peter Bannister, Michael Brady, and Stephen Smith. 2002. “Improved Optimization for the Robust and Accurate Linear Registration and Motion Correction of Brain Images.” NeuroImage 17 (2):825–41.

Magnotta, Vincent A., Lee Friedman, and FIRST BIRN. 2006. “Measurement of Signal-to-Noise and Contrast-to-Noise in the fBIRN Multicenter Imaging Study.” Journal of Digital Imaging 19 (2):140–47.

Markov, N. T., M. M. Ercsey-Ravasz, A. R. Ribeiro Gomes, C. Lamy, L. Magrou, J. Vezoli, P. Misery, et al. 2014. “A Weighted and Directed Interareal Connectivity Matrix for Macaque Cerebral Cortex.” Cerebral Cortex 24 (1). Oxford University Press:17–36.

Mars, Rogier B., Saad Jbabdi, Jérôme Sallet, Jill X. O’Reilly, Paula L. Croxson, Etienne Olivier, Maryann P. Noonan, et al. 2011. “Diffusion-Weighted Imaging Tractography-Based Parcellation of the Human Parietal Cortex and Comparison with Human and Macaque Resting-State Functional Connectivity.” The Journal of Neuroscience: The Official Journal of the Society for Neuroscience 31 (11):4087–4100.

McLaren, Donald G., Kristopher J. Kosmatka, Terrance R. Oakes, Christopher D. Kroenke, Steven G. Kohama, John A. Matochik, Don K. Ingram, and Sterling C. Johnson. 2009. “A Population-Average MRI-Based Atlas Collection of the Rhesus Macaque.” NeuroImage 45 (1):52–59.

Mennes, Maarten, Bharat B. Biswal, F. Xavier Castellanos, and Michael P. Milham. 2013. “Making Data Sharing Work: The FCP/INDI Experience.” NeuroImage 82:683–91.

Miranda-Dominguez, Oscar, Brian D. Mills, David Grayson, Andrew Woodall, Kathleen A. Grant, Christopher D. Kroenke, and Damien A. Fair. 2014. “Bridging the Gap between the Human and Macaque Connectome: A Quantitative Comparison of Global Interspecies Structure-Function Relationships and Network Topology.” The Journal of Neuroscience: The Official Journal of the Society for Neuroscience 34 (16):5552–63.

Mortamet, Bénédicte, Matt A. Bernstein, Clifford R. Jack Jr, Jeffrey L. Gunter, Chadwick Ward, Paula J. Britson, Reto Meuli, Jean-Philippe Thiran, Gunnar Krueger, and Alzheimer’s Disease Neuroimaging Initiative. 2009. “Automatic Quality Assessment in Structural Brain Magnetic Resonance Imaging.” Magnetic Resonance in Medicine: Official Journal of the Society of Magnetic Resonance in Medicine / Society of Magnetic Resonance in Medicine 62 (2):365–72.

Paxinos, George. 2009. The Rhesus Monkey Brain in Stereotaxic Coordinates. Academic Press.

Paxinos, George, Xu-Feng Huang, and Arthur W. Toga. 1999. The Rhesus Monkey Brain in Stereotaxic Coordinates. Academic Press.

Power, Jonathan D. 2017. “A Simple but Useful Way to Assess fMRI Scan Qualities.” NeuroImage 154 (July):150–58.

Reveley, Colin, Audrunas Gruslys, Frank Q. Ye, Daniel Glen, Jason Samaha, Brian E Russ, Ziad Saad, Anil K Seth, David A. Leopold, and Kadharbatcha S. Saleem. 2017. “Three-Dimensional Digital Template Atlas of the Macaque Brain.” Cerebral Cortex 27 (9):4463–77.

Rilling, J. K. 2014. “Comparative Primate Neuroimaging: Insights into Human Brain Evolution.” Trends in Cognitive Sciences 18 (1). Elsevier Current Trends:46–55.

R. Matthew Hutchison, Stefan Everling. 2012. “Monkey in the Middle: Why Non-Human Primates Are Needed to Bridge the Gap in Resting-State Investigations.” Frontiers in Neuroanatomy 6. Frontiers Media SA. https://doi.org/10.3389/fnana.2012.00029

Rohlfing, Torsten, Christopher D. Kroenke, Edith V. Sullivan, Mark F. Dubach, Douglas M. Bowden, Kathleen A. Grant, and Adolf Pfefferbaum. 2012. “The INIA19 Template and NeuroMaps Atlas for Primate Brain Image Parcellation and Spatial Normalization.” Frontiers in Neuroinformatics 6 (December):27.

Saleem, Kadharbatcha S., and Nikos K. Logothetis. 2012. A Combined MRI and Histology Atlas of the Rhesus Monkey Brain in Stereotaxic Coordinates. Academic Press.

Seidlitz, Jakob, Caleb Sponheim, Daniel Glen, Frank Q. Ye, Kadharbatcha S. Saleem, David A. Leopold, Leslie Ungerleider, and Adam Messinger. 2017. “A Population MRI Brain Template and Analysis Tools for the Macaque.” NeuroImage, April. https://doi.org/10.1016/j.neuroimage.2017.04.063.

Seidlitz, Jakob, František Váša, Maxwell Shinn, Rafael Romero-Garcia, Kirstie J. Whitaker, Petra E. Vértes, Paul Kirkpatrick Reardon, et al. 2017. “Morphometric Similarity Networks Detect Microscale Cortical Organisation And Predict Inter-Individual Cognitive Variation.” https://doi.org/10.1101/135855.

Shmuel, A., and D. A. Leopold. n.d. “Neuronal Correlates of Spontaneous Fluctuations in fMRI Signals in Monkey Visual Cortex: Implications for Functional Connectivity at Rest. - PubMed - NCBI.” Accessed November 21, 2017. https://www.ncbi.nlm.nih.gov/pubmed/18465799.

“The Average Baboon Brain: MRI Templates and Tissue Probability Maps from 89 Individuals.” 2016. NeuroImage 132 (May). Academic Press:526–33.

Vanderwal, Tamara, Clare Kelly, Jeffrey Eilbott, Linda C. Mayes, and F. Xavier Castellanos. 2015. “Inscapes : A Movie Paradigm to Improve Compliance in Functional Magnetic Resonance Imaging.” NeuroImage 122:222–32.

Vanduffel W, Et al. n.d. “Monkey Cortex through fMRI Glasses. - PubMed - NCBI.” Accessed November 21, 2017. https://www.ncbi.nlm.nih.gov/pubmed/25102559.

Van Essen, David C. 2002. “Windows on the Brain: The Emerging Role of Atlases and Databases in Neuroscience.” Current Opinion in Neurobiology 12 (5):574–79.

Van Essen, David C. 2012. “Cortical Cartography and Caret Software.” NeuroImage 62 (2):757–64.

Van Essen, David C., Matthew F. Glasser, Donna L. Dierker, and John Harwell. 2012. “Cortical Parcellations of the Macaque Monkey Analyzed on Surface-Based Atlases.” Cerebral Cortex 22 (10):2227–40.

Van Essen, D. C., and M. F. Glasser. n.d. “In Vivo Architectonics: A Cortico-Centric Perspective. - PubMed - NCBI.” Accessed November 21, 2017. https://www.ncbi.nlm.nih.gov/pubmed/23648963.

Vincent, J. L., G. H. Patel, M. D. Fox, A. Z. Snyder, J. T. Baker, D. C. Van Essen, J. M. Zempel, L. H. Snyder, M. Corbetta, and M. E. Raichle. 2007. “Intrinsic Functional Architecture in the Anaesthetized Monkey Brain.” Nature 447 (7140):83–86.

Xu, Ting, Arnaud Falchier, Elinor Sullivan, Gary Linn, Julian Ramirez, Deborah Ross, Eric Feczko, et al. 2017. “Delineating the Macroscale Areal Organization of the Macaque Cortex in Vivo.” bioRxiv. https://doi.org/10.1101/155952.

Zhang D, Et al. n.d. “Diffusion Tensor Imaging Reveals Evolution of Primate Brain Architectures. - PubMed - NCBI.” Accessed November 21, 2017. https://www.ncbi.nlm.nih.gov/pubmed/23135357.

